# Designing Robustness to Temperature in a Feedforward Loop Circuit

**DOI:** 10.1101/000091

**Authors:** Shaunak Sen, Jongmin Kim, Richard M. Murray

**Affiliations:** The Department of Electrical Engineering, Indian Institute of Technology Delhi, Hauz Khas, New Delhi 110016, INDIA.; The Division of Biology and Biological Engineering (BBE), California Institute of Technology, Pasadena, CA 91125, USA.; The Divisions of Engineering and Applied Science (EAS) and Biology and Biological Engineering (BBE), California Institute of Technology, Pasadena, CA 91125, USA.

## Abstract

‘Incoherent feedforward loops’ represent important biomolecular circuit elements capable of a rich set of dynamic behavior including adaptation and pulsed responses. Temperature can modulate some of these properties through its effect on the underlying reaction rate parameters. It is generally unclear how to design such a circuit where the properties are robust to variations in temperature. Here, we address this issue using a combination of tools from control and dynamical systems theory as well as preliminary experimental measurements towards such a design. We formalize temperature as an uncertainty acting on system dynamics, exploring both structured and unstructured uncertainty representations. Next, we analyze a standard incoherent feedforward loop circuit, noting mechanisms that intrinsically confer temperature robustness to some of its properties. Further, we explore different negative feedback configurations that can enhance the robustness to temperature. Finally, we find that the response of an incoherent feedforward loop circuit in cells can change with temperature. These results present groundwork for the design of a temperature-robust incoherent feedforward loop circuit.

## I. INTRODUCTION

Living cells are subject to a wide variety of environmental changes, for example, due to seasonal changes in variables like temperature and humidity. The biomolecular circuits that regulate their behavior are likely to possess the property that ensures they function robustly in face of such environmental changes [1]. A similar property can be desirable for engineered biomolecular circuit designs, whereby they also function robustly in a range of environments [2]. This can have multiple benefits, including promoting their efficacy in a wider range of environments and allowing circuit modules designed in slightly different environments to be reliably interconnected. In particular, temperature is an important environmental variable from the point of view of biomolecular circuit design. This is because temperature can modulate the functional output of biomolecular circuit designs as a direct consequence of its effect on the underlying reaction rate parameters. Indeed, for biomolecular circuit designs, temperature has been identified as a key element of the environmental context against which functional robustness needs to be assessed [2].

Insofar as temperature robustness in biomolecular circuits is concerned, the most studied systems have been limit cycle oscillators, both natural [3] and synthetic [4]. In these systems, temperature robustness manifests in the oscillation period, which is approximately constant over a range of temperatures. One way of characterizing the robustness has been to define a temperature coefficient 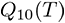 of a temperature-dependent rate/property *k*(*T*), as the ratio 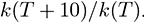 A *Q*_10_ = 2 means that *k* doubles for a 10°C increase in temperature. In contrast, a *Q*_10_ = 1 means that *k* does not change with temperature. It is often found that *Q*_10_ of the oscillation period is close to 1, indicating temperature robustness, against the typical values in the range 2–3 for biomolecular reaction rates [5]. A recent example where temperature robustness has also been studied is bacterial chemotaxis [6]. In these examples, a key way in which temperature robustness is achieved is for two temperature-dependent rates to exactly cancel the effects of each other so that the functional output is temperature-robust. These studies provide important early work towards the design of temperature-robust biomolecular circuits.

Feedforward loop circuits are a class of biomolecular circuits with at least three striking features [7]. First is their widespread occurrence in different biomolecular contexts, underscored by their overrepresentation in transcriptional networks [8]. Second is their systems level properties, for example, the incoherent feedforward loop can exhibit both perfect adaptation [9], [10] and fold-change detection [11], [12]. Third is the dynamical outputs that can be achieved, for example, the incoherent feedforward loop can generate pulse dynamics, whose quantitative properties such as the pulse height, adaptation value, rise time, and decay time can be of functional importance in cells. What are the different ways in which temperature can affect these properties (Fig. 1)? Can we design modifications to an incoherent feedforward loop circuit that can enhance their temperature robustness? Indeed, it is generally unclear how to design an incoherent feedforward loop circuit where properties are temperature-robust.

**Fig. 1.**
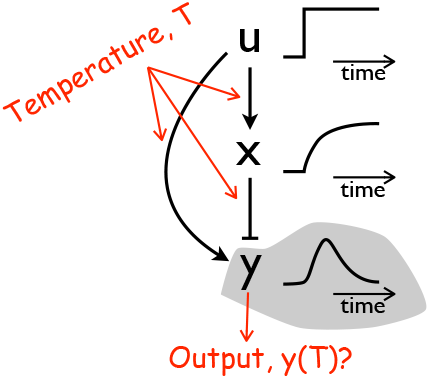
Temperature can affect the output of an incoherent feedforward loop through its effect on the underlying reactions. A standard representation of such a circuit is sketched. In response to an input step, there can be a pulsed output. Red arrows indicate the propagation of temperature dependence.

Here, we ask the question of how to design a temperature-robust incoherent feedforward loop circuit. To address this issue, we use tools from control and dynamical systems theory. We model the effect of temperature as both a structured and an unstructured uncertainty acting on a biomolecular circuit. Next, we investigate a standard incoherent feedforward loop circuit model and note inherent features that promote temperature robustness. We also investigate design variants using negative transcriptional feedback with the aim of enhancing this robustness. Finally, as a step towards a robust design, we present preliminary experimental data illustrating how the dynamics of a circuit realization change with temperature. These results should aid the design of a temperature-robust incoherent feedforward loop circuit.

## II. RELEVANT THEORETICAL TOOLS

In order to understand how temperature affects functional output in biomolecular circuits and how this effect may be compensated for, we first adapt relevant tools from control and dynamical systems theory in this context. Consider the mathematical representation of the dynamics of a biomolecular circuit obtained using standard mass-action-based ordinary differential equations,

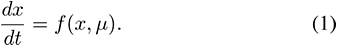

Here, *x* is a vector of concentration variables and *μ* is a vector of reaction rate parameters which depend on temperature *T*. Because of the temperature dependence of reaction rate parameters 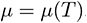, for example, as characterized by a temperature coefficient 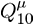 in the range 2–3, functional properties of this system such as steady-state levels or transient features can also be temperature dependent.

In fact, using this characterization, effect of temperature in Equation (1) can be analyzed like a structured uncertainty in the reaction rate parameters. For example, the range of *Q*_10_’s of the outputs under consideration can be computed for a given set of *Q*_10_’s of the reaction rate parameters. Alternatively, these parametric uncertainties map to a range of values for each output under consideration, which can be directly considered. In particular, the temperature dependence of the steady-state values *x*_0_ can be obtained directly from the algebraic equation *f*(*x*_0_, *µ*) = 0.

In order to investigate other potentially useful uncertainty representations, we consider a linearization of Equation (1) around the operating point *x* = *x*_0_,

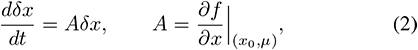

where *δx* = *x* − *x*_0_ is the small variation in the state variables. Each element of the matrix 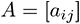 will exhibit some variation due to a temperature, corresponding to a variation in the value of one or more parameters. Suppose in a temperature range, 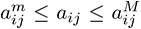. Consequently, we can express the uncertainty in each element as,

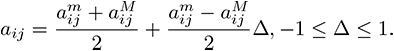

Equivalently, using 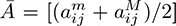 and 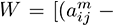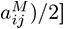, we can express 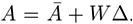 Then, the transfer function from the initial condition *x*_0_ to the state *x* is (*sI* − 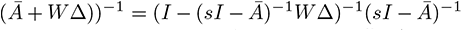 (Fig. 2). We note that this is analogous to a feedback uncertainty representation around the nominal system 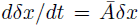, with the uncertainty *W*Δ itself being static in nature [13].

**Fig. 2.**
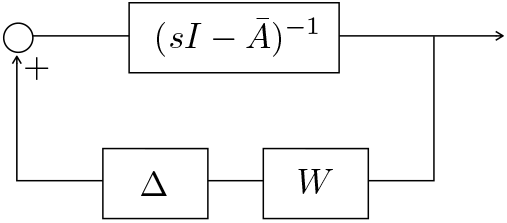
Effect of temperature can be modeled as an unstructured feedback uncertainty.

For example, consider a simple model of protein production-degradation,

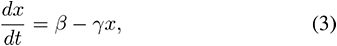

where *x* is the concentration of a protein, produced at a constant rate *β*, and degraded as a first-order process with rate constant *γ*. At steady state, 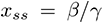. Clearly, the temperature dependence in this model can be analyzed by directly considering the parametric uncertainty in *β* and *γ* due to temperature. Additionally, in considering the linearization about *x* = *x_ss_*, *dδx*/*dt* = −*γδx*, we can obtain the unstructured feedback uncertainty representation for *γ*^*m*^ ≤ 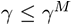 with 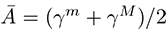 and 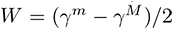. Such uncertainty representations can be useful in assessing the robustness of certain properties or controller designs that exist for the nominal system.

In order that perfect temperature compensation be achieved, a natural question arises with regard to the properties that such a controller must possess. In fact, this question bears resemblance to the Internal Model Principle [13], [14], [15], which requires that the system possesses a model of the external signal that it needs to be robust against. For example, perfect adaptation to step inputs can be implemented through the use of an integrator block in the loop transfer function. If the input was a ramp, the presence of a single integrator block would not suffice for perfect adaptation. In this case, the requirement is for two integrator blocks.

To check whether a similar guideline is relevant for the problem of perfect temperature compensation, where it is desired for an output to be invariant to temperature even though underlying circuit parameters change with temperature, we augmented the simple model above with a control input *u* that is assumed to be able to directly control the rate of change of *x*,

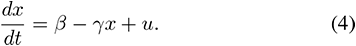

We consider the steady state *x_ss_* as the output of this system. For simplicity, we assume that it is only *β* that changes with temperature, *β* = *β*(*T*). This assumption serves to localize the effect of temperature on one reaction term, helping with this analogy to perfect adaptation. In this context, perfect temperature compensation can be achieved if the function *β* = *β*(*T*) is known. A control input that achieves this is 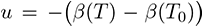, where *T*_0_ is a reference temperature. Therefore, knowledge of temperature dependence and such feedback cancellation can enable perfect temperature compensation.

An adaption of these tools in the context of temperature robustness illustrates a way to analyze temperature robustness and the ideal conditions for a system response to be independent of temperature. In the following sections, we focus on a standard incoherent feedforward loop circuit model where these tools are relevant.

## III. INHERENT TEMPERATURE ROBUSTNESS IN AN INCOHERENT FEEDFORWARD LOOP MODEL

To characterize possible temperature dependence of the properties of an incoherent feedforward loop, we consider a standard mathematical model of it (adapted from [12]),

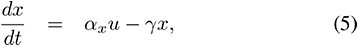

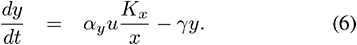

Here, *u* is the input, *x* is the intermediate variable, and *y* is the output. The parameters *α*_*x*_, *α*_*y*_, *γ*, and *K_x_* represent the production rate of *x*, of *y*, their dilution rates, and the binding constant of *x* to the promoter of *y*, respectively. The origin of the term *K_x_*/*x* is as an approximation to a repression function like *K_x_*/(*K_x_*+*x*). We assume that the default values of the parameters are *α*_*x*_ = *α*_*y*_ = *γ* = *K_x_* = *u* = 1. Further, we assume that a step change in the input leads to a change in the value of *u* from 1 to 2. This model exhibits the adaptation property (Fig. 3): In response to a step change in input *u*, both *x* and *y* first increase; As *x* increases further, it represses *y* and *y* relaxes to its pre-step value. Therefore, the *y* waveform exhibits a pulse in response to a step change in input *u* (Fig. 3). This pulse shape can be further characterized by the following four properties: adaptation level (*y_eq_*), pulse amplitude (*y_m_*), rise time (*τ_τ_*), and decay time (*τ_d_*). Each of these features can be temperature dependent. This is because they depend on reaction rate parameters which can themselves be temperature dependent. The above model presents a medium to investigate the possible temperature dependencies.

**Fig. 3.**
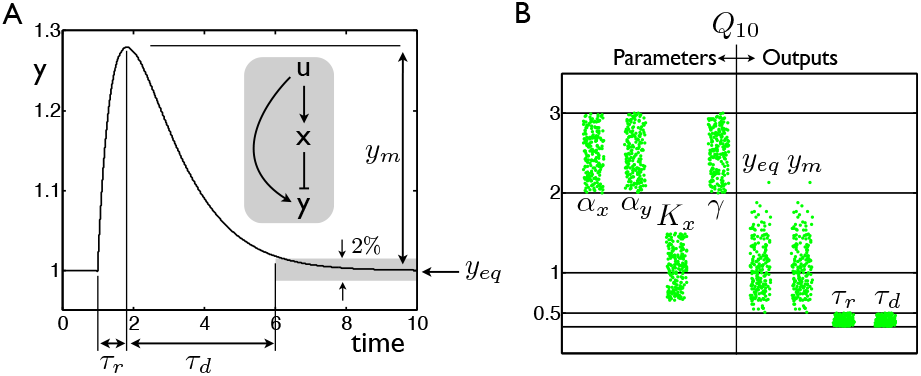
Propagation of temperature dependence in a model of an incoherent feedforward loop circuit. A. Black line is a trajectory computed from the model. Different properties are graphically illustrated on the trajectory. Inset shows a schematic illustration of the circuit. B. Green dots illustrate the range of *Q*_10_’s of the indicated reaction rate parameters and outputs. Horizontal spread is due to an arbitrary random number added for illustration purposes. Solid black lines indicate key *Q*_10_ values.

We note that the adaptation property is solely dependent on the mechanism and robust to changes in parameters. As such, it is also robust to parametric variations owing their origin to a temperature effect. In contrast, properties of the pulsed response, such as adaptation level, pulse amplitude, rise time, and decay time can depend on temperature through their dependence on circuit parameters. To investigate this dependence in the model, we make the following assumptions. First, we assume that the parameters take the default values specified above. Second, we assume that their temperature coefficient *Q*_10_ is in the range 2–3. Finally, based on this assumed range of *Q*_10_’s of the parameters, we calculate the range of temperature coefficient *Q*_10_ of each of the properties. This approach treats temperature as a structured parameter uncertainty and assesses the robustness when temperature is increased by 10°C.

As an illustration of this approach, consider the equilibrium value of 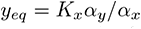 obtained in the above model. In particular, let us consider the ratio *r* = **α**_*y*_/**α**_*x*_. Assuming that the *Q*_10_’s of **α**_*x*_ and **α**_*y*_ are in the range 2–3, the *Q*_10_ of *r* has a minimum value of 2/3 ≈ 0.66 and a maximum value of 3/2 = 1.5. As the DNA binding constant *K_x_* arises as a ratio of the off and on rates for the binding of *X* to promoter of gene *y*, we assume that its *Q*_10_ is also in the range 0.66–1.5 (assuming these off/on rates also have *Q*_10_ in range 2–3). Therefore, we estimate the *Q*_10_ of *y_eq_* to be in the range 0.44–2.25. In fact, this illustration captures a temperature compensation mechanism already inherent in this model. The temperature coefficient of the equilibrium level is different from those of the parameters because the parameters combine as a ratio of terms. Whenever two terms with similar temperature dependencies combine in a ratio, the effective temperature dependence of the ratio can be attenuated. Indeed, in this case, the production rates **α*_x_* and **α*_y_* are expected to have similar temperature dependencies, even if they do not strictly fall in the range of *Q*_10_’s assumed above. Therefore, this temperature compensation is expected to persist in a general setting as well and is an instance of the feedback cancellation mentioned in the previous section.

Next, we apply this approach to other properties. As obtaining their analytical expressions is not as straightforward as that for the equilibrium value, we resort to numerical simulations. For this, we first choose *Q*_10_ of each of the parameter as a different random number in the range 2–3. Using this selection, we compute the *Q*_10_ for each of the above-mentioned properties: Pulse amplitude is computed as the maximum of the resulting waveform away from the equilibrium value. The rise time is computed as the time taken to reach the maximum pulse amplitude from when the pulse is applied. Similarly, the decay time is computed as the time it takes to go from when the pulse maximum is reached to when the response first returns to within 2% of its equilibrium value. The results of this computation for *N* = 200 different random choices of *Q*_10_ sets are shown in Fig. 3. As expected, we find that the *Q*_10_ of the equilibrium value is in the range 0.44–2.25. Similarly, the pulse amplitude is also in this range, indicating a similar compensation effect to a change in temperature. In contrast, the *Q*_10_’s of the rise time and decay time are in the range 0.33–0.5, consistent with timescale being determined by the reciprocal of *γ*, a parameter with *Q*_10_ in the range 2–3. Therefore, this indicates that other than a conversion of a 100%–200% increase in parameters to a 50%–66% reduction in the property, there is no other temperature robustness effect for the timescale-related parameters.

Finally, an implicit source of temperature robustness in the model is the supposition that both *x* and *y* are effectively degraded via dilution in the same manner. If their effective degradation rates are dissimilar, say *γ*_*x*_ and *γ*_*y*_ for *x* and *y* respectively, then a possible difference in how they change with temperature may lead to an added temperature sensitivity in the properties. For example, the equilibrium value *y_eq_* = *K_x_*α*_y_γ*_*x*_/(**α**_*x*_*γ*_*y*_) indicates that the temperature coefficient *Q*_10_ lies in the range 0.3–3.4, which is an expanded version of the range estimated above. Therefore, the modeling assumption that *x* and *y* have the same degradation constant provides an inherent cancellation of temperature dependencies.

To summarize, this standard incoherent feedforward loop circuit model already has three inherent features that can enhance the temperature robustness of the pulse height and the equilibrium value. These three are,

1. Similar temperature dependencies of the production rates.
2. Proportionality to the DNA binding constant, which is itself a ratio of two rates.
3. Same effective degradation terms acting on *x* and *y*.

## IV. DESIGNS FOR TEMPERATURE ROBUSTNESS

With the aim of enhancing robustness of these properties to temperature, we explore modifications to this circuit. Motivated by the presence of negative feedback in numerous robustness contexts, we explored the effect of adding negative transcriptional feedback to the circuit.

### 1. Negative feedback of *y* on itself

Consider a model, similar to above, where *y* negatively feeds back onto itself,

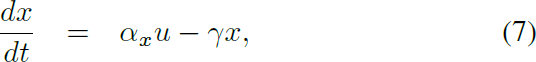

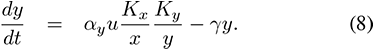

We find that the adaptation property can persist in this model (see Fig. 4A). In fact, the equilibrium value, *y_eq_* = 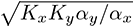. In comparing this value with the one obtained for the above model, we note the presence of a square root. This square root can be significant from the point of view of temperature robustness. If a rate k has a *Q*_10_ in the range 2–3, then 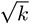 has a *Q*_10_ in the range 1.41–1.73, which indicates enhanced temperature robustness. Consistent with this, we find that the *Q*_10_ of *y_eq_* should lie in the range 0.54–1.84. This is a slightly narrower range than that obtained in the model without transcriptional feedback. Indeed, this is also seen when the *Q*_10_ ’s of parameters are chosen randomly (Fig. 4B). These computations show a slight enhancement in the temperature robustness properties of the adaptation value as well as the peak pulse amplitude. For these parameters, there is no significant difference in how the timescale properties of the pulse depend on temperature between this model and the main model. In general, the presence of the square root can have an effect of reducing the temperature dependence compared to the main model even if the production rates and binding constants have other temperature dependencies different from the ones assumed here.

**Fig. 4.**
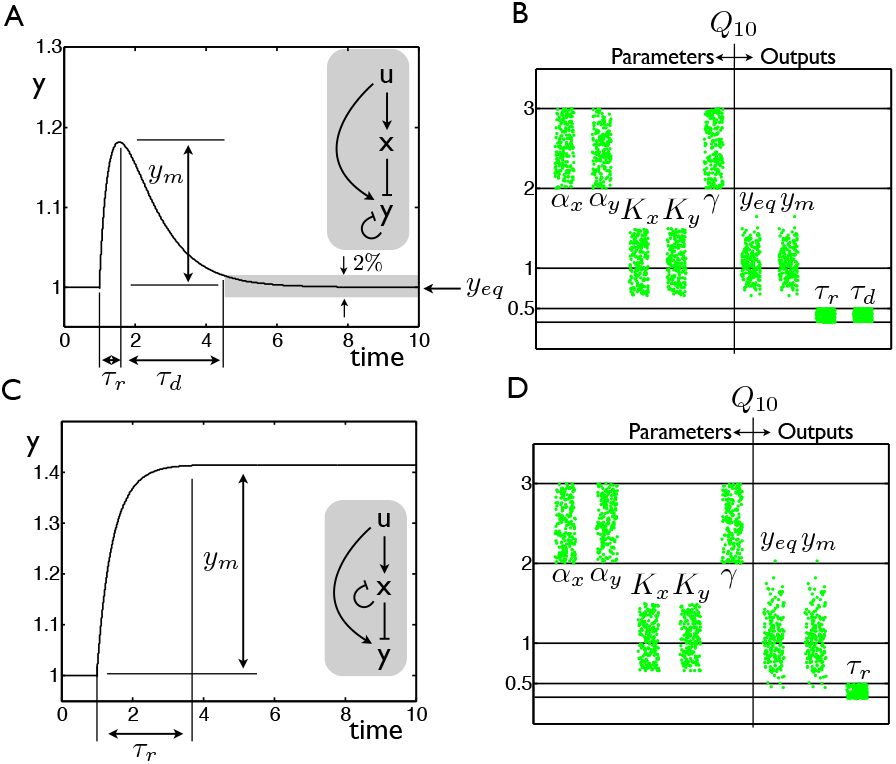
Effect of different negative feedback configurations on the propagation of temperature dependence in an incoherent feedforward loop circuit. A–B. Negative feedback is added from *y* onto itself. C–D. Negative feedback is added from *x* onto itself. Simulations are performed as noted previously. Additional parameters *K_y_ = J_x_* = 1.

### 2. Negative feedback of *x* on itself

Next, we consider the case where *x* negatively feeds back on itself. The corresponding model is,

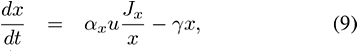

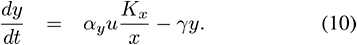

Here, we find that the equilibrium value itself depends on the input, 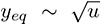. This dependence on the input indicates that the adaptation property itself does not exist. Nevertheless, we performed a similar analysis as above to gauge how temperature may affect output properties (Fig. 4C–D). We find that the output amplitudes corresponding to the maximum and final values, *y_m_* and *y_eq_*, respectively, have a larger spread relative to temperature, whereas the timescale *τ_R_* depends on temperature similar to the above cases.

These two models illustrate the effect of adding different negative feedback configurations with the aim of modifying temperature robustness of the pulse property of the incoherent feedforward loop circuit. Of the two, we find that the negative feedback from *y* onto itself can help enhance temperature robustness of the amplitude properties.

## V. PRELIMINARY EXPERIMENTS

As a step towards the experimental design of an incoherent feedforward loop circuit that is temperature-robust, we performed a preliminary experiment measuring the circuit dynamics of such a circuit inside cells at different temperatures. The goal of this experiment was to observe whether or not temperature actually affects the behavior of an actual circuit realization inside growing cells. This implementation of an incoherent feedforward loop uses transcriptional interactions (Fig. 5A): Transcription factor AraC can activate the expression of the transcriptional repressor TetR and the green fluorescent protein deGFP. Both proteins are expressed under the AraC-activable *P_bad_* promoter. Additionally, TetR binding sites are inserted into the *P_bad_* promoter controlling expression of deGFP so that it is repressible by TetR. An advantage of using the transcription factors AraC and TetR is that their activity can be modulated using inducers arabinose and anhydrotetracycline (aTc), respectively. Finally, AraC is expressed from a constitutive promoter that is regulated by the housekeeping sigma factor *σ*^70^. We realized that this promoter was temperature-sensitive in that its expression increases with increasing temperature. Further, this sensitivity is a consequence of the activity of a mutant cI protein, which can additionally affect the growth rate. Nevertheless, we proceeded with measurements of this circuit to get a sense of whether and how temperature can affect the response.

**Fig. 5.**
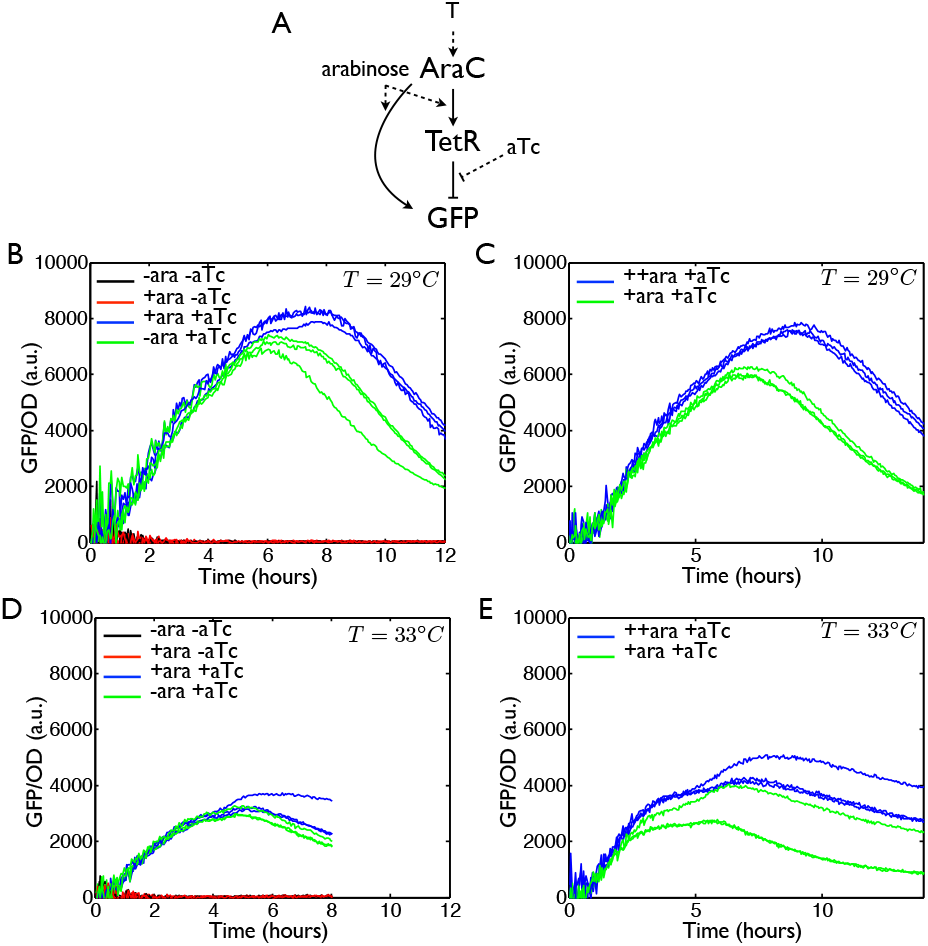
Preliminary experimental measurements characterizing the temperature dependence of an incoherent feedforward loop circuit realization. A. Schematic illustration of the circuit realization. B. and C. are measurements at 29° C D. and E. are measurements at 33°C. Each measurement is performed in triplicate on the same day. Further, the raw fluorescence and raw OD values were background subtracted using well with only media and no cells (autofluorescence was found to be similar to this blank fluorescence in this media). In each panel, different colors represent different inducer combinations. ++ara denotes 0.018% arabinose. +ara denotes 0.0018% arabinose. +aTc denotes 0.001 *µ*/ml aTc. -ara and -aTc denote the absence of these inducers.

Measurements of circuit dynamics were performed in a platereader (BioTek Synergy H1) capable of measuring both fluorescence and optical density of cells over time and at different temperatures. For this, a single colony from an LB plate was picked and grown overnight in a clear, rich media (MOPS-glycerol, Teknova M2105 with 50% glycerol instead of 20% glucose) at 29°C. This liquid culture was diluted 1:50 into fresh media and grown for a second overnight cycle at the same temperature. Then, the culture was diluted 1:50 in fresh media and grown for around 2 hours at the same temperature. This culture was used for assays with appropriate amounts of inducers. Measurements were performed in a 96-well plate sealed with a Breathe-EZ membrane. The plate reader protocol alternated double-orbital shaking with measurements of the optical density (600 nm, denoted OD) and green fluorescent protein (excitation 485 nm, emission 525 nm) at 29°C, with measurements performed at 3 minute intervals. All growth steps are performed with appropriate antibiotics.

We found that the circuit could exhibit a pulse response (Fig. 5B). Interestingly, this pulse response depended on the presence of the inducer aTc. Indeed, the circuit exhibited no pulse in the absence of aTc. This is likely due to the strong repressing effect of TetR. Additionally, there was a small increase in the height of the pulse as arabinose levels were increased. This was consistent with the expectation from the circuit diagram. To check this further, two different arabinose levels were used in the presence of aTc (Fig. 5C). Again, the circuit behaved as expected in that the pulse height increased for a higher arabinose level. Overall, we found an expected pulse-like shape for this circuit, but the apparent dominant effect of aTc needed to be investigated further.

To investigate whether this response is temperature dependent, we repeated the above protocol with the platereader measurement being performed at 33°C (Fig. 5D,E). The circuit exhibited a pulse-like shape at this temperature as well, similar to that observed at 29°C. However, we noted that the maximum pulse amplitude as well as the time taken to reach this amplitude were different than at 29°C. These results provide initial evidence for the dependence of both the pulse height and timescale of the output of this circuit design on temperature. We note the two non-idealities of this circuit realization due to the apparent dominant effect of the inducer aTc and the temperature-sensitive promoter expressing AraC. Indeed, addressing the latter through the use of a different promoter can directly reduce the effect of temperature on both promoter strengths and the growth rate, corresponding to the model parameters **α*_x_*, **α*_y_* and *γ*, respectively. Therefore, these measurements naturally raise questions of how to make this design more temperature-robust, by addressing these non-idealities as well as through the models analyzed in previous sections.

## VI. CONCLUSIONS AND FUTURE WORK

The design of biomolecular circuit devices whose function is robust to temperature is a key challenge. Here, we used tools from control and dynamical systems theory to guide the goal of designing a temperature-robust pulse-generating incoherent feedforward loop circuit, presented as the following results. First, we adapt these tools to present structured and unstructured uncertainty representations modeling the role of temperature in biomolecular circuits. Second, we computationally study a standard incoherent feedforward loop circuit, pointing out inherent circuit features that promote robust performance. Third, we studied design modifications using negative transcriptional feedback, finding that negative transcriptional feedback at the output stage can further improve this robustness to temperature. Fourth, we present preliminary experimental results showing that the response of an initial design for an incoherent feedforward loop circuit can change with temperature. These results lay the groundwork for the design of a temperature-robust incoherent feedforward loop circuit.

In the context of achieving temperature robustness, an interesting aspect is how much knowledge about the temperature dependence of the circuit parameters is required. From one point of view, there are mechanisms that can promote temperature robustness without information of the temperature dependence of circuit parameters *per se*. An example presented here where this occurs is in the first design modification, where addition of a square root expression can reduce the *Q*_10_ of the output. On the other hand, it is reasonable to expect knowledge of the temperature dependence of the circuit parameters to aid the design of temperature compensation. As presented through a simple production-degradation example and also in the inherent robustness features of the feedforward loop circuit, this may also aid in cancelling the effect of temperature through an appropriate feedback term. This situation seems similar to the Internal Model Principle, where the presence of an internal model of a disturbance in a system allows it to completely attenuate the effect of the disturbance.

A natural extension of this study is to proceed with the construction of variant incoherent feedforward loop circuit designs that exhibit different predictable temperature-robustness properties. In addition to experiments in cells, the use of a cell extract transcription-translation environment may also be helpful by providing a faster iterative design cycles [16]. As part of this, the first step is to construct an alternative to the basic incoherent feedforward loop circuit to verify the predicted inherent temperature robustness properties. A second step is to add the negative transcriptional feedback at the output stage and check for the predicted enhancement in robustness. Additionally, it should motivate the development of mathematical tools for uncertainty representations as design aids.

Investigating the design of temperature-robust biomolecular circuits is part of a larger goal of designing biomolecular circuits that robustly function in a range of environments. Through these results, we have presented foundational steps towards the design of a temperature-robust incoherent feedforward loop, a biomolecular circuit that is both widespread and exhibits a rich set of qualitative and quantitative dynamic behavior. These results should help in the design of other temperature-robust circuits as well as to further analyze temperature-robustness in naturally occurring biomolecular circuits.

## ACKNOWLEDGMENT

We gratefully acknowledge S. Guo and C. Hayes for their help with the experimental part of this work, especially for providing the *E. coli* strain containing the biomolecular circuit used and for the measurement protocol. Research supported in part by the Gordon and Betty Moore Foundation and the NSF Molecular Programming Project.

